# Comprehensive chemotyping, and the gonadal regulation, of seven kisspeptinergic neuronal populations in the mouse brain

**DOI:** 10.1101/2024.07.23.604881

**Authors:** Vito S. Hernández, Mario A. Zetter, Oscar R. Hernández-Pérez, Rafael Hernández-González, Ignacio Camacho-Arroyo, Robert P. Millar, Lee E. Eiden, Limei Zhang

## Abstract

**Background:** Kisspeptinergic signaling is well-established as crucial for regulation of reproduction, but its potential broader role in brain function is less understood. This study investigates the distribution and chemotyping of kisspeptin-expressing neurons within the mouse brain.

**Methods:** RNAscope singleplex, duplex and multiplex *in situ* hybridization methods were used to assess kisspeptin mRNA (*Kiss1)* expression and its co-expression with other neuropeptides, excitatory and inhibitory neurotransmitter markers, and sex steroid receptors in intact and gonadectomized young adult mice.

**Results:** Seven distinct kisspeptin neuronal chemotypes were characterized, including within two novel *Kiss1*-expressing groups described here for the first time: the ventral premammillary nucleus, and the nucleus of the solitary tract. *Kiss1* mRNA was also localized in the soma, and within the dendritic compartment, of hypothalamic neurons. Altered *Kiss1* expression following gonadectomy suggests a previously unappreciated role for androgen receptors in regulating kisspeptin signaling.

**Conclusion:** This study provides a detailed chemoanatomical map of kisspeptin-expressing neurons in the brain, highlighting their potential functional diversity. The discovery of new kisspeptin-expressing neuronal populations, and gonadectomy-induced changes in *Kiss1* expression patterns, provide a basis for further exploration of non-endocrine roles for kisspeptin in brain function.

## 1. Introduction

The kisspeptins, which include the precursor KP-54, and the proteolytically processed KP-14, KP-13, and KP-10 ^(1, 2)^, are neuropeptides first identified in human melanoma cells as products of the metastasis-suppressor gene *Kiss1* ^(3)^. Kisspeptin (KP) and its receptor (deorphanized from GPR54, KiSS1R) ^(4)^, are important for cell signaling within the hypothalamus regulating reproductive function through stimulation of gonadotropin-releasing hormone (GnRH) secretion ^(5-14)^. KP neurons are also located outside of the hypothalamus, notably in the amygdala, where emotion and cognition are integrated and putatively linked to reproduction and reproductive behaviors ^(15-19)^.

Early *in situ* hybridization (ISH) histochemistry and immunohistochemistry in mice revealed five kisspeptin mRNA (*Kiss1*)-expressing neuronal clusters in the mouse brain ^(5, 6, 20-25)^; the most prominent being in the arcuate nucleus (Arc), and in the rostroperiventricular region of the third ventricle (RP3V). This region includes the periventricular nucleus (PeN), the anteroventral periventricular nucleus (AVPV), and some scattered cells in the anterodorsal preoptic area. The other regions reported to express *Kiss1*, albeit at lower levels, are the medial amygdala, the bed nucleus of the stria terminalis and the dorsomedial hypothalamus ^(26)^. Although both sexes express *Kiss1* mRNA in all five areas, expression in the AVPV has been reported to be sexually dimorphic, with higher expression of *Kiss1* mRNA and KP in females than in males ^(5, 22)^.

The regulation of KP expression and release is complex and may be brain region-specific, both during development, and as a result of epigenetic and environmental factors such as sex steroids, gonadal status, stress and nutrient state ^(27-32)^. Gonadal status and KP signaling mutually regulate the hypothalamic-pituitary-gonadal (HPG) axis.

KP neurons express other neuropeptides that contribute to the dynamics of GnRH regulation ^(13)^. KP^Arc^ neurons express kisspeptin, neurokinin B, and dynorphin (and are dubbed “KNDy” neurons for this reason) ^(33-35)^ as well as galanin ^(36)^. Neurokinin B and dynorphin act autosynaptically on KP neurons in the Arc to synchronize and regulate the pulsatile secretion of kisspeptin and corresponding release of GnRH from fibers in the median eminence ^(13)^.

KP *per se* plays an excitatory neurotransmitter role acting upon transient receptor potential canonical (TRPC) and potassium channels expressed in a subpopulation of GnRH neurons ^(37, 38)^. TRPC channels are a subfamily of nonselective cation channels that are part of the TRP superfamily ^(39, 40)^. They are calcium-permeable and receptor-operated, and are activated by the phospholipase C (PLC) signaling pathway ^(38)^. Effects on GnRH excitability exerted by KP neurons which co-express either gamma-aminobutyric acid (GABA) or glutamate have been studied mainly in the KP^RP3V^ and KP^Arc^ populations, respectively ^(41) (42)^. KP’s direct excitatory effects on GnRH neurons can be potently modulated by GABA, via GABA_A_, and GABA_B_, receptors, and by glutamate, in a spatial-temporal and steroid-dependent manner ^(12-14, 43)^. Kisspeptins can also activate non-GnRH neurons that synapse upon GnRH neurons, exerting indirect effects on GnRH neuronal responses by modulating fast synaptic transmission through GABA and glutamate ^(44) (45)^. As a result of this complex regulation, many circuit-level questions remain about the pathways, direct and indirect, through which KP cooperates with amino acid and neuropeptide co-transmitters to control GnRH neurons, as well as non-HPG neuronal activity controlling various behaviors. Detailed chemotyping of KP neurons throughout the brain will be helpful in answering these questions.

The objectives of our study were first, to characterize kisspeptinergic neuronal populations, based on their molecular signatures and distribution throughout the mouse brain using single, dual, and multiplex RNAscope methods and second, to assess the impact of gonadectomy on each of these KP neuronal populations in both male and female mice. Seven distinct *Kiss1-*expressing populations and their sensitivity to gonadectomy are described in detail, including two novel KP-expressing groups, *i*.*e*., the *Kiss1* populations in the ventral premammillary nucleus, and in the nucleus of solitary tract. *Kiss1* mRNA was also observed in both soma and dendritic compartments of hypothalamic neurons. Altered *Kiss1* expression occurs following gonadectomy (GNX) in male mice, suggesting a previously unappreciated role for androgen receptors in regulating KP signaling. The chemoanatomical mapping of KP neurons throughout the brain provides a foundation for a better understanding of how KP neurons modulate sexual reproduction and potentially other reproductive behavior-related sensory-emotion-cognitive functions through their peptidergic, glutamatergic and GABAergic synaptic connections.

## 2. Materials and methods

### 2.1. Animals

C57BL/6 young adult mice of 25-30 g (10-12 weeks old) were obtained from the local animal vivarium, distributed in four groups (N=24, n=6, males vs. females and intact vs. GNX), and housed three per cage under controlled temperature and illumination (12h/12h) with water and food *ad libitum*. Mice subjected to GNX were singly-housed for recovery for one week after surgery and then returned to their home cages. All animal procedures were approved by the local research ethics supervision committees (license UNAM: CIEFM-079-2020, and NIMH-IRP ACUC).

### 2.2. Gonadectomy (GNX)

The detailed procedures of mouse ovariectomy (OVX) and orchietectomy (ORX) are described elsewhere ^(46)^. The absence of vaginal proestrus smears was confirmed by performing vaginal cytology during consecutive days post-surgery.

### 2.3 Assessment of reproductive cycle

Vaginal smears, followed by the method described elsewhere ^(47, 48)^, were obtained from intact female subjects during consecutive days and proestrus stage was chosen to collect brains of mice. Successful GNX was confirmed post-mortem.

### 2.4 RNAscope single, dual, and multiplex *in situ* hybridization (ISH, DISH, MISH) procedures

Detailed methods are described elsewhere ^(49-51)^. Briefly, after 7-8 weeks of gonadectomy male and female mice were deeply anaesthetized with sodium pentobarbital (100 mg/kg b.w., i.p.) and decapitated using a small animal guillotine. Brains from intact and gonadectomized mice were removed and rapidly frozen in pulverized dry ice. The fresh-frozen brains were sectioned in sagittal and coronal planes, of 12 μm thickness, with a Leica CM-1520 cryostat and mounted on positively charged microscope slides. Every fourth section was processed. Experimental procedures were performed according to manufacturer instructions for <u>RNAscope™ 2.5 HD Assay - BROWN</u>.

To evaluate additional mRNAs in *Kiss1*-positive neurons relevant to potential co-release of additional peptides and neurotransmitters, and sex steroid responsiveness, in the seven populations of identified *Kiss1* expressing neurons, we performed DISH and MISH experiments according to manufacturer instructions (acdbio.com) with the following probes: KP (Mm-*Kiss1*, Cat No. 500141), VGAT (Mm-Slc32a1-C2, Cat No. 319191-C2), VGLUT2 (Mm-Slc17a6-C2, Cat No. 428871-C2); neurokinin B (Mm-Tac2-C2, Cat No. 446391-C2), dynorphin (Mm-Pdyn-C2, Cat No. 318771-C2), estrogen receptor alpha (Mm-Esr1-C2, Cat No. 496221-C2), androgen receptor (Mm-Ar-C2, Cat No. 316991-C2), PACAP (Mm-Adcyap1*-C2*, Cat No. 405911-C2) and neurotensin (Mm-Nts-C3, Cat No. 420441-C3).

### 2.5 Imaging and analysis

To quantify the expression level of *Kiss1* in the Singleplex (brown) ISH-reacted sections, micrographs were taken in a given region of interest (ROI) chosen based on consistent anatomical landmarks across samples and experimental groups. Micrographs (>5) were taken per region/subject. In each micrograph a region of 0.04 mm^2^ area (200 μm × 200 μm), within the ROI was delimited for analysis.

For co-expression semiquantitative analysis we scored the colocalization, at cellular level i.e., around or over the same Nissl-labeled nucleus, with the following criteria: “n.o.”, not observed; “+” for 1 in 5 cells with *Kiss1* labeling; “++” 2 in 5 cells; “+++” 3 in 5, and “++++” 4 in 5 (or more).

To assess the GNX effect on *Kiss1* expression, two measurements were performed. For the two *Kiss1*-expressing neuronal populations in which dendritic *Kiss1* labeling was observed, we used the criterion of cells with *Kiss1* dendritic expression vs total *Kiss1* labeled cells in the 0.04 mm^2^ area within Arc or RP3V. For the rest of the regions, we counted the *Kiss1* labeling dots over the Nissl-labeled nuclei and made average per region per experimental condition, in the 0.04 mm^2^ area (for more consideration, see results section with focused discussion).

### 2.6 Statistical analysis

Statistical analysis was performed using GraphPad Prism 10 Software. Data were assessed for normality using D’Agostino-Pearson test. To compare the fraction of *Kiss1*-positive neurons with dendritic mRNA expression, or the average number of *Kiss1*-positive mRNA puncta between groups for each of these regions, a two-way ANOVA with sex and treatment as factors was used. A Tukey post-hoc test was used for pairwise comparisons. Differences were considered statistically significant at *p* < 0.05. Different letters above the bars in the graphs were used to indicate significant differences between groups.

## 3. Results

### 3.1 Kisspeptin neuronal group in the rostro-periventricular area of the hypothalamus (KP^RP3V^)

The RP3V region (Fig. 1A) hosts one of the two major *Kiss1-*expressing neuronal groups, which has been extensively studied over the past two decades. It encompasses *Kiss1* neurons distributed through the ventromedial preoptic nucleus (VMPO), the anteroventral periventricular nucleus (AVPV), the periventricular nucleus (Pe), and the median preoptic nucleus (MnPO). The density of *Kiss1*-expressing neurons is heterogeneous. In a parasagittal view (Fig. 1, A1), around lateral 0.05 mm, heterogeneously distributed *Kiss1-*labeled cells can be observed extending around 300 μm in the rostro-caudal dimension and 800 μm in ventro-dorsal dimension, with the density gradient decreasing upward. In the median preoptic nucleus, sparsely distributed *Kiss1* expressing neurons can be found (Fig. 1, A1a). In contrast, in a coronal view (Fig. 1A2, bregma 0.02 mm), *Kiss1* labeled cells are located in rather narrow bands alongside the third ventricle wall, with most of the cells falling into a less than 100 μm band (each side) and the density decreasing sharply in the medio-lateral dimension (Fig. 1A2a).

**Figure 1.**
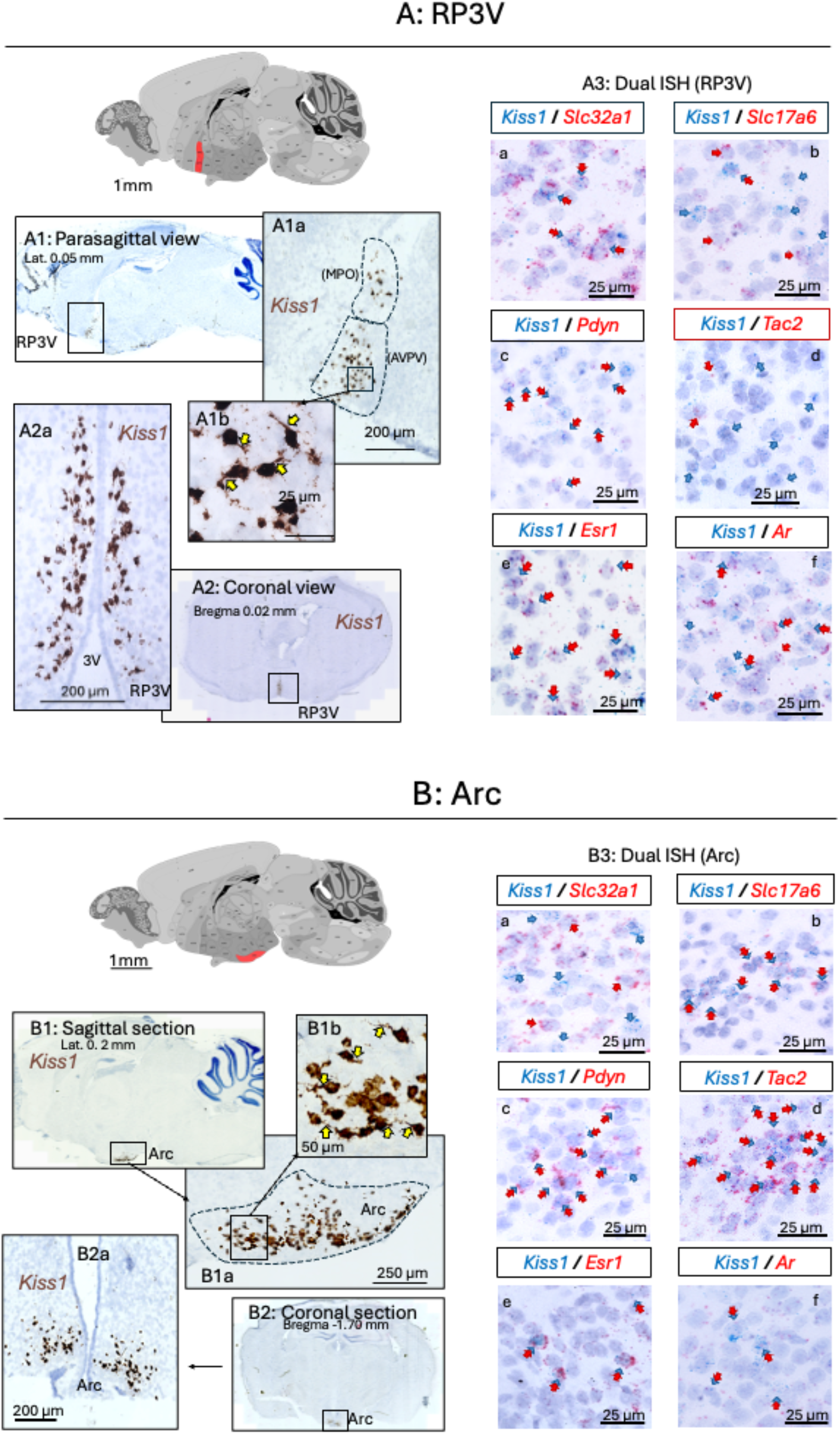
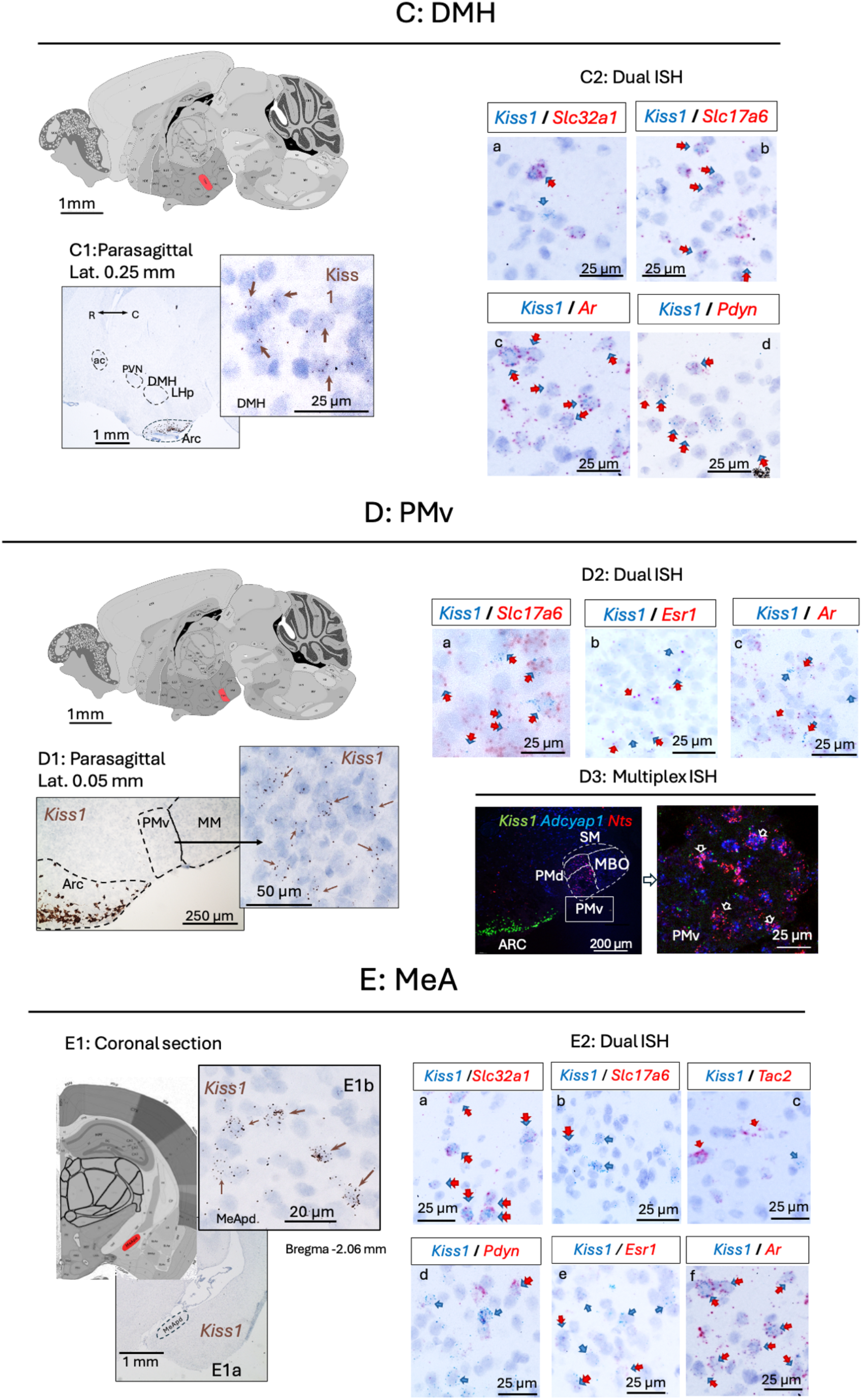
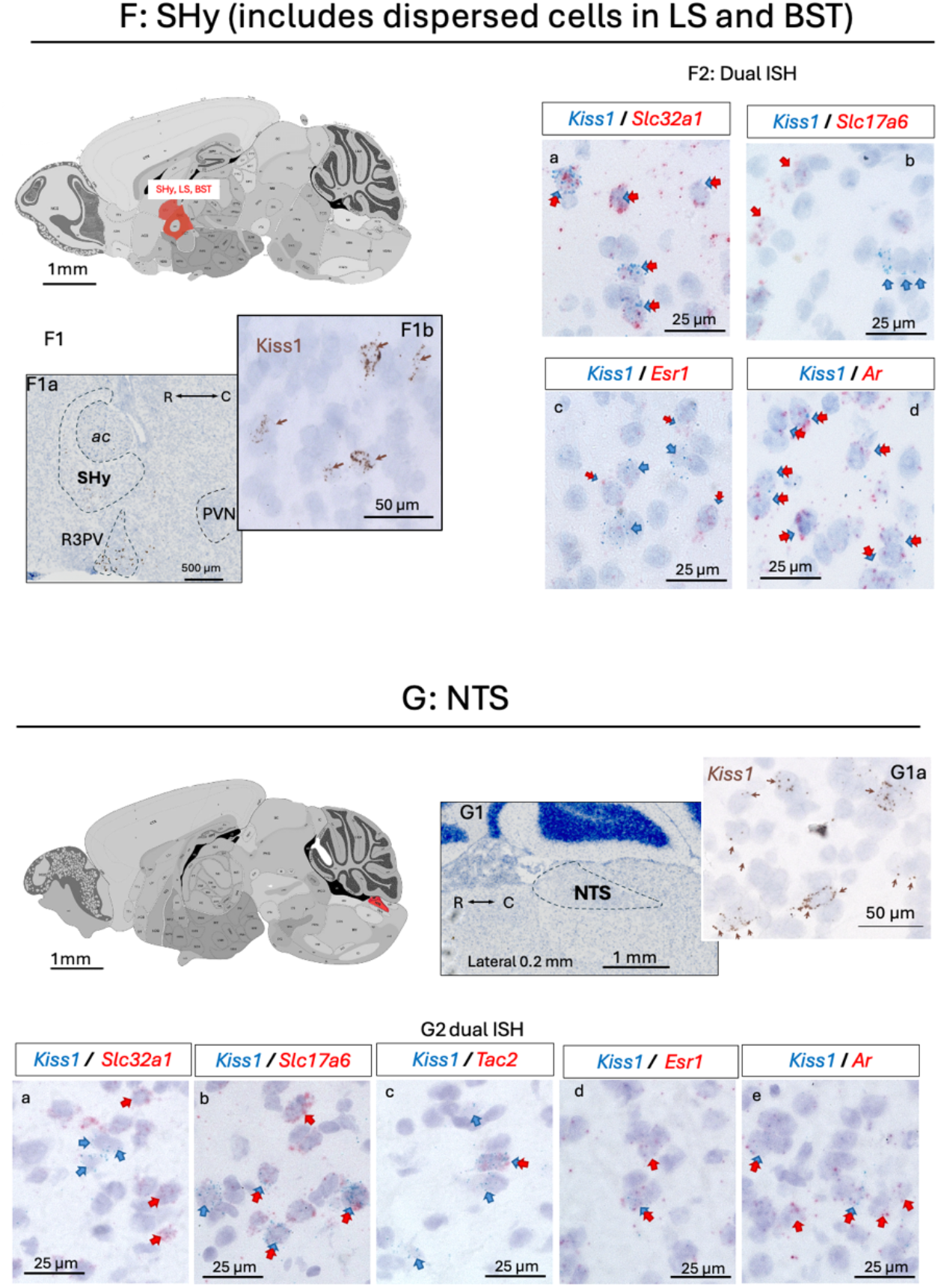
Anatomical localization and chemotyping of seven Kiss1-expressing neuronal populations in the mouse brain. A: KP population in the rostro-periventricular area of the hypothalamus (KP^RP3V^) shown in a parasagital (A1 and inset) and in a coronal view (A2) have a heterogeneous spatial distribution in the rostro-caudal axis and a steep decrease in density in the medial to lateral direction. The yellow arrows in the figures indicate cells with prominent expression of Kiss1 mRNA in the dendritic compartment. A_3_, panels show RNAscope duplex reactions that demonstrate the expression of Kiss1 (blue arrows), and other mRNAs (red arrows) coding for Slc32a1, Slc17a6, Tac2, Pdyn, Esr1 or Ar. Double arrows show that in this region Kiss1 is co-expressed with Slc32a1, Pdyn, Esr1 and Ar. B: Kisspeptin population in the arcuate nucleus of the hypothalamus (KP^ARC^). B1b and inset show Kiss1 expressing neurons with dendritic Kiss1 expression (yellow arrows). B3 panels show that in this population Kiss1 is colocalized with Slc17a6, Tac2, Pdyn, Esr1 and Ar mRNAs but not with Slc32a1. C: Kiss1 population in the dorso-medial hypothalamic area (KP^DMH^). Kiss1 detected by singleplex RNAscope is shown in a parasagittal section in C1, Kiss1-expressing neurons in the DMH are indicated by brown arrows in the inset. Duplex RNAscope micrographs in C2 show that in this population Kiss1 is colocalized with Slc17a6, Slc32a1, Pdyn and AR. No co-expression with Tac2, or Esr1 mRNAs was observed. D: Kiss1 expressing neuronal population in the ventral premammillary nucleus area (KP^PMv^). D1 and inset: micrograph of the PMv, taken from a sagittal section shows Kiss1 epressing neurons scattered in the PMv, the expression levels of Kiss1 was low. DISH (D2) and MISH (D3) RNAscope reactions show that this newly characterized population of Kiss1 neurons co-express the mRNAs for Slc17a6 (D2a), Esr1 (D2b), AR (D2c), PACAP, and neurotensin (Adcyap1 and Nts respectively, D3). E: Kiss1 expressing population in the medial amygdala (KP^MeA^). Panel E1, low and high magnification micrographs of a coronal section showing Kiss1 expressing neurons concentrated in the posterodorsal part of the medial amygdala (MeApd). E2, Duplex RNAscope showed that Kiss1 neurons strongly co-expressed Slc32a1(E2a) and Ar (E2f) and to a lower level also expressed Slc17a6 (E2b), Esr1 (E2e), and Pdyn (E2d). F: Kiss1 expressing population in the septo-hypothalamic area (KP^SHy^). Panel F1, parasagittal section depicts the SHy area, spanning septal, BST and hypothalamic regions where scattered low Kiss1-expressing neurons (inset of F1) were found around the anterior commisure. F2, micrographs taken from SHy in slices treated for RNAscope Duplex. Kiss1 was observed to be colocalized with Slc32a1 (F2a), Esr1 (F2c) and Ar (F2d). No co-expression with Tac2, Pdyn or Slc17a6 (F2b) mRNAs was observed in this region. G: Kiss1 expressing population in the NTS (KP^NTS^). Singleplex RNAscope (G1), showed Kiss1 expressing neurons located in the solitary tract nucleus of male and female mice. G1 and G1a, photomicrograph of a sagittal section depicting neurons expressing Kiss1 at low abundance. G2 panels show by RNAscope Duplex the co-expression of Kiss1 mRNA with Slc17a6 (G2b), Tac2 (G2c), Esr1 (G2d) and Ar (G2e), no co-expression with Slc32a1 (G2a) or Pdyn was observed. The performance of each assay was validated by examining other regions within the same brain slice known to express the second mRNA, ensuring that the absence of colocalization was not due to technical issues but rather accurately reflected the absence of co-expression in the specific neuronal population.

*Kiss1* is extensively expressed in proximal dendrites of KP^RP3V^ population, in addition to cell bodies (Fig. 1 inset, yellow arrows). This phenomenon has been interpreted to suggest local dendritic translation of mRNA. Dendritic *Kiss1* expression was observed in both male and female brain.

The RNAscope DISH method allows evaluation of the co-expression of separate mRNAs within single cells (Fig. 1A3). This *Kiss1* neuronal population co-expresses *Slc32a1* (mRNA for VGAT, Fig. 1A3a) and mostly without *Slc17a6* (mRNA for VGLUT2, Fig. 1A3b) expression. Some *Kiss1* neurons express *Pdyn* (mRNA for dynorphin Fig. 1A3c) but are negative for *Tac2* (mRNA for neurokinin B, Fig. 1A3d). The KP^RP3V^ group strongly co-expressed *Esr1* and *Ar* (mRNAs encoding the estrogen receptor alpha and androgen receptor respectively, Fig. 1A3e and Fig. 1A3f, respectively, double blue/red arrows indicate co-expression), suggesting that the KP neurons in the KP^RP3V^ regions are sensitive to estrogens and androgens acting via ERα and AR.

### 3.2 Kisspeptin neuronal group in the arcuate hypothalamic area (KP^Arc^)

The hypothalamic arcuate nucleus hosts a second extensively studied hypothalamic KP-expressing group of neurons (Fig. 1B, a sagittal view of KP^Arc^ at lateral 0.25 mm). These cells also showed both somatic and dendritic expression of *Kiss1* (Fig. 1B1a and 1B1b, yellow arrows). The cells are densely distributed toward the ventral floor of the arcuate nucleus with a rostro-caudal span of around 1 mm (Fig. 1B1a). The density decreases dorsally (Fig. 1B1a). In Fig. 1B1a and Fig. 1B2a, the *Kiss1* population is shown from a coronal view at bregma -1.70 mm, in a spherical shape with a diameter of approximately 250 μm on each side of the third ventricle. Using DISH (Fig. 1B3), we confirmed the co-expression of mRNAs for the vesicular glutamate transporter 2 (*Slc17a6*, Fig. 1B3b), neuropeptides neurokinin B (*Tac2*, Fig. 1B3d) and dynorphin (*Pdyn*, Fig. 1B3c), as previously reported ^(33, 35, 52)^. *Kiss1*-positive neurons strongly expressed estrogen receptor alpha (*Esr1* Fig. 1B3e) and weakly expressed androgen receptor (*Ar*, Fig. 1B3f) mRNAs. KP^Arc^ neurons did not co-express *Slc32a1*, the mRNA encoding VGAT (Fig. 1B3a).

### 3.3 Kisspeptin neuronal group in the dorso-medial hypothalamic area (KP^DMH^)

*Kiss1*-expressing neurons were found in the dorso-medial hypothalamic region (DMH, Fig. 1C), beginning immediately caudal to the paraventricular nucleus posterior division and extending to the lateral posterior hypothalamic region (LHp, Fig. 1C1). The cells of this group had low *Kiss1* expression (Fig. 1C1 inset). Analysis of DISH experiments (Fig. 1C2) showed that about half of the KP^DMH^ cells express *Slc17a6 (*Fig. 1C2b), and some express *Slc32a1 (*Fig. 1C2a), *Ar (*Fig. 1C2c) and *Pdyn (*Fig. 1C2d), but not *Esr1* or *Tac2* mRNAs (micrographs not shown).

### 3.4 Kisspeptin neuronal group in the ventral premammillary nucleus (KP^PMv^)

A newly described population of cells expressing KP mRNA (*Kiss1*) was identified caudally to the Arc, in the ventral premammillary nucleus (PMv, Fig 1D). These neurons had a low expression of KP (average 5-6 puncta per cell) but were densely packed and homogeneously distributed throughout the PMv (Fig. 1D1 inset). Using dual and multiplex *in situ* hybridization (Fig. 1D2) we showed that these *Kiss1*+ neurons co-express the vesicular glutamate transporter 2 mRNA (*Slc17a6*, Fig. 1D2a) and show a low expression of *Ar* and *Esr1* (Fig. 1D2b and c). They also express the mRNA for PACAP (*Adcyap1*) and neurotensin (*Nts*) (Fig. 1D3). No co-expression of *Tac2* or *Pdyn*, mRNAs that are strongly expressed in KP neurons of the adjacent arcuate region, was observed.

### 3.5 *Kiss1*-expressing neuronal populations in medial amygdala (KP^MeA^)

A population of *Kiss1*-expressing neurons exists in the medial postero-dorsal nucleus of the amygdala (KP^MeA^, Fig 1E and E1a and E1b). This extrahypothalamic population has been studied by several groups ^(17, 19, 53)^. The amygdalar *Kiss1* neurons co-expressed *Slc32a1* (mRNA for VGAT, Fig. 1E2a), with a few expressing VGLUT2 (*Slc17a6*, Fig. 1E2b). Some of these neurons co-express dynorphin (*Pdyn*, Fig. 1E2d), but not neurokinin B (*Tac2*, Fig. 1E2c). KP^MeA^ neurons co-expressed *Ar* and, at a lower level *Esr1* (Fig. 1E2e and E2f).

### 3.6 *Kiss1*-expressing neuronal populations in septo-hypothalamic nucleus (KP^SHy^) and surrounding area of the anterior commissure extending to lateral septum (LS) and bed nucleus of stria terminalis (BST)

Dispersed *Kiss1*-expressing neurons with molecular signatures similar to those of the medial amygdala group *(KP*^*MeA*^) were found surrounding the caudal part of the anterior commissure, spanning the septo-hypothalamic (SHy) area ^(54)^, the lateral septum (LS) and the bed nucleus of stria terminalis (Fig. 1F and F1a and F1b), confirming previous reports using IHC ^(55)^, ISH and marker protein expression in transgenic animals ^(24, 56, 57)^. These neurons co-express *Slc32a1* (Fig. 1F2a), as well as *Esr1* and *Ar* (Fig.1 F2c and F2d). No *Slc17a6* (Fig. 1F2b), *Pdyn* or *Tac2* were not detected in these neurons (micrograph not shown).

### 3.7 *Kiss1*-expressing neuronal populations in the nucleus of tactus solitarii (NTS)

The metencephalic tractus solitarii (NTS) contains the most caudal population of KP neurons, previously reported based on metastin immunoreactivity ^(58, 59)^. Here, we show the expression of kisspeptin mRNA (Fig. 1G1 and 1G1a) in NTS. Analysis of DISH experiments (Fig. 1G2) showed that KP neurons expressed VGLUT2 (*Slc17a6*, Fig. 1G2b), with approximately 40% of them expressing *Tac2* (Fig. 1G2c), 25% *Esr1* (Fig. 1G2d), and most expressing *Ar* (Fig. 1G2e). No co-expression of *Pdyn* or *Slc32a1* (Fig. 1G2a) was observed.

### 3.8 *Kiss1* expression’s vulnerability of the seven populations by gonadectomy

Seven weeks after GNX, there was an evident reduction in *Kiss1* expression levels in the KP^RP3V^ and an increase in the KP^Arc^ cell populations observed under the microscope on a per-cell average basis. However, given the high sensitivity of the RNAscope assay and the heterogeneous distribution of this population of *Kiss1* neurons, we found that assessing changes in density of cells with increased *Kiss1*-labeled dots, in a given section, was an insufficiently quantitative criterion to reflect the observed reduction/augmentation. We therefore sought a new method to assess the changes in *Kiss1* expression after short-term GNX, specifically by quantifying the percentage of *Kiss1*-positive neurons that expressed mRNA in the dendritic compartment.

The *Kiss1* neurons of female proestrous mice were more strongly labelled, and a significantly higher (*p*<0.001) percentage of them showed dendritic mRNA expression, compared to males (71.2% ± 3.5% in females vs. 27.5% ± 2.6% in males), see Fig. 2A and A1 vs. A3. After GNX there was a reduction in the number of cells expressing dendritic *Kiss1*: OVX (5.1 % ± 1.5%, p< 0.001 vs. female intact) and ORX (ORX: 1.7% ± 1.2%, p< 0.001 vs. male intact).

**Figure 2.**
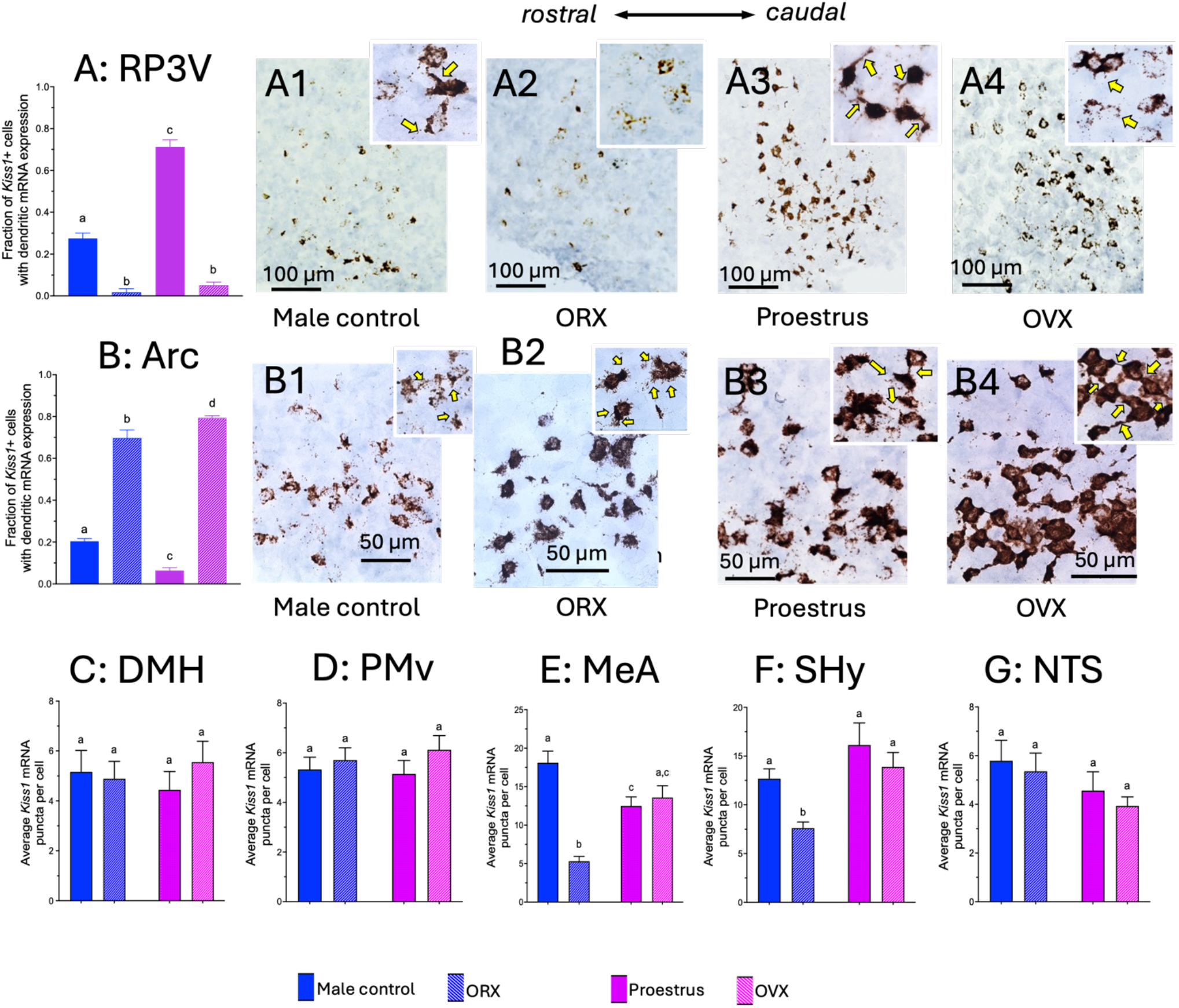
Effect of gonadectomy on Kiss1 expression in the seven Kiss1 expressing populations of the mouse brain. A-B: The effects of gonadectomy were quantified in the two major population of Kiss1 expressing neurons, namely the RP3V (panels As) and Arc (panels Bs). The neurons in these regions showed a high cellular expression of Kiss1, and a heterogeneous spatial distribution. These two populations were distinguished by the expression of Kiss1 in the dendritic compartment. Thus, we measured the fraction of Kiss1 cells that displayed dendritic Kiss1 expression in relation to the total number of cells expressing Kiss1. This allows a more precise quantification of the effects of gonadectomy, minimizing possible errors derived from the high sensitivity of the test that is able to detect single molecules of mRNA thus detecting cells with insignificant expression and also sampling errors derived from the very heterogeneous spatial distribution of the cells, particularly in the RP3V. Representative photomicrographs of RP3V (A1-A4) and Arc (B1-B4) show low and high magnification of these areas in male and female proestrus control mice and after orchidectomy (ORX) and ovariectomy (OVX). Yellow arrows indicate examples of neurons with dendritic expression. In the RP3V the fraction of cells expressing dendritic Kiss1 is higher in proestrus female than in control males, and GNX significantly reduced the dendritic expression in both sexes. In the Arc the male control animals showed a higher fraction of Kiss1 cells with dendritic expression, and after GNX the Kiss1 expression was dramatically increased in both sexes, with the OVX females showing a higher fraction of Kiss1 dendritic expression than the ORX males. Significant (p< 0.05) statistical differences between groups are indicated by letters. Bars without shared letters are significantly different. C-F: The expression of Kiss1 in these neuronal populations was much lower than in the RP3V and Arc. For these regions we quantified the average mRNA puncta per cell. No significant differences in Kiss1 expression levels between males and females, nor any effect after gonadectomy was detected in the hypothalamic KP^DMH^ (panel C) and KP^PMv^ (panel D). For the extrahypothalamic regions, in the medial amygdala (KP^MeA^, panel E) and the septo-hypothalamic region (KP^SHy^, panel F) we observed that gonadectomy caused a significant reduction in Kiss1 expression only in the male animals. For the nucleus of tractus solitarius population (KP^NTS^, panel G), no differences in Kiss1 expression between males and females or after gonadectomy were observed.

In the KP^Arc^ region the quantification of the percentage of KP^ARC^ neurons with somatodendritic *Kiss1*-expression versus total labeled cells (Fig. 2B) showed a significantly higher percentage of dendritic *Kiss1*-expressing neurons in intact male mice (20.4% ±1.24%) compared to female proestrus mice (6.4% ±1.4 %). After GNX, there was a potent upregulation of *Kiss1* dendritic expression in the Arc region in both sexes, with the expression levels in females surpassing that of males (ORX: 69.7% ± 3.86% vs OVX: 79.3% ± 1.02%, p < 0.05).

Quantification of the average number of *Kiss1* mRNA puncta per cell in the KP^DMH^, KP^PMv^ and KP^NTS^ showed no differences between male and female mice, or any effect of GNX in males or females (Fig. 2C, 2D, 2G).

In KP^MeA^, GNX induced a significant decrease in the average number of *Kiss1* mRNA puncta per cell only in male mice (Fig 2E). Similarly to the *KP*^*MeA*^ population, only male animals showed a decrease in the expression of *Kiss1* after GNX in *KP*^*SHy*^ neurons, from 12.68 ± 1.0 to 7.62 ± 0.62 *Kiss1* puncta per cell (Fig. 2 F).

An overview of the seven populations of neurons in the mouse brain, with regard to the coexpression of KP mRNA (*Kiss1*) with *Slc32a1* or *Slc17a6*, is represented in the drawing of Fig. 3A, in which the regions containing mainly *Kiss1* neurons co-expressing *Slc32a1* are represented in red, the regions with *Kiss1* neurons mainly co-expressing *Slc17a6* are represented in green, and the DH region containing both *Slc32a1* and *Slc17a6 Kiss1* neurons is represented in blue. With the exception of the MeA (indicated with a dotted line border), all the regions are located very close to the midline. Fig. 3B, summarizes the molecular signatures of the seven populations of kisspeptinegic neurons, indicating the co-localization of *Kiss1* with *Slc32a1, Slc17a6, Pdyn, Tac2, Esr1* and *Ar*. A semiquantitative report on the co-expression ratio of these mRNAs with *Kiss1* is provided. Also a summary of the effect of ORX and OVX on *Kiss1* levels for each of the regions described in this study is presented in Fig. 3B.

**Figure 3.**
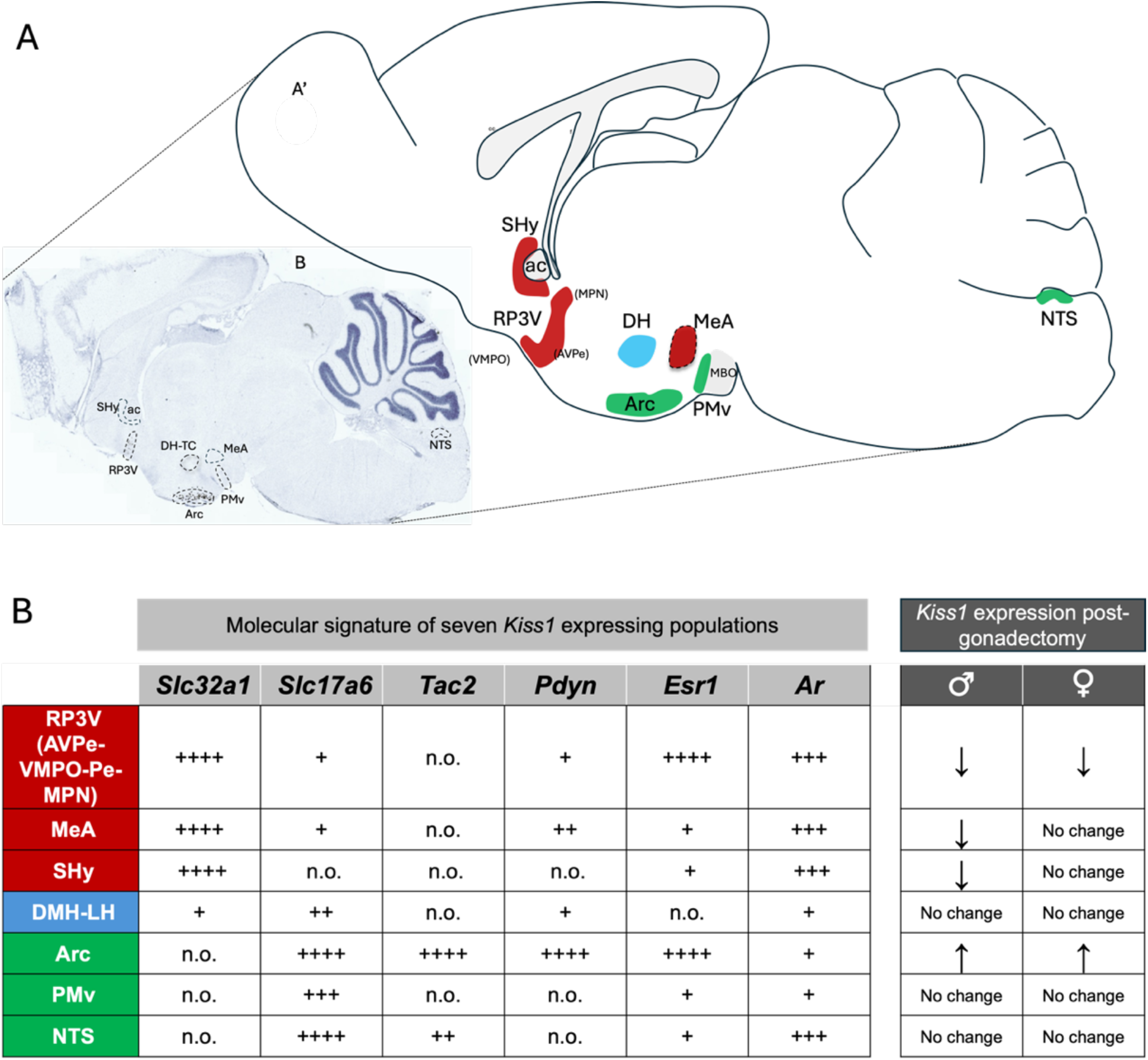
Molecular signature of seven population of Kiss1 neurons in the brain, A: parasagittal section and representative drawing of the regions that were observed to host Kiss1 positive neurons. The regions that have mainly GABAergic (Slc32a1) are colored in red, the regions with mainly glutamatergic (Slc17a7) are colored in green, the DM region colored in blue, contains glutamatergic and GABAergic Kiss1 neurons. B: Summary table depicting the co-expression of Kiss1 mRNA with the mRNAs coding for the vesicular GABA transporter (VGAT, Slc32a1); the vesicular glutamate transporter 2 (VGLUT2, Slc17a6); neurokinin B (Tac2), Dynorphin (Pdyn), estrogen receptor alpha (Esr1) and androgen receptor (Ar). The semiquantitative scores represent percentages of co-expression with Kiss1 neurons: (n.o.) for not observed, (+) for <25%, (++) for 26-50%, (+++) for 51-75% and (++++) for >75% Abbreviations: anterior commissure (ac); anteroventral periventricular nucleus (AVPe); arcuate nucleus (Arc); dorsomedial hypothalamic nucleus (DMH); mammillary bodies (MBO); medial amygdala (MeA), median preoptic nucleus (MPO), nucleus of tractus solitarius (NTS); septohypothalamic area (SHy); rostro-periventricular nu. of the third ventricle (RP3V); ventral premammillary nucleus (PMV); ventromedial preoptic area (VMPO).

## 4. Discussion

### 4.1 Physiological implications of the seven KP mRNA-expressing populations

i. We report here the molecular signatures of seven distinct *Kiss1*-expressing populations and their sensitivity to gonadectomy. For the two major KP hypothalamic groups, namely the rostro-periventricular group (KP^RP3V^) and the arcuate hypothalamic group (KP^Arc^), our observations mainly concur with reports in the literature. However, using the highly sensitive “Brown kit” RNAscope method, we observed, and report here for the first time, that in these two populations, besides *Kiss1* mRNA expression in the soma, a high proportion of KP cells express mRNA in *the dendritic compartment*. This phenomenon suggests an ability to express and translate KP mRNA in dendrites ^(60) (61-63)^.

The RP3V KP neurons play a key role in the positive feedback of estrogen, which is critical for the stimulation of gonadotropin-releasing hormone (GnRH) resulting in the preovulatory surge of luteinizing hormone (LH) which culminates in ovulation. Notably, these neurons express estrogen receptor alpha (ERα), and their activation enhances *Kiss1* expression, which subsequently stimulates GnRH neurons to release GnRH in a pulsatile manner ^(64-66)^. We demonstrated that the KP^RP3V^ population co-expresses *Slc32a1, Esr1, Ar* and *Pdyn* RNAs. In agreement with the majority of previous reports characterizing mRNA expression in this region we observed sexual dimorphism and the effects of GNX ^(20, 23, 67, 68)^. We observed a higher number of KP-expressing neurons in females than in males and a down-regulation of the density of *Kiss1*-expressing neurons in both sexes after GNX. This down-regulation of *Kiss1* expression after gonadectomy was particularly significant in the dendritic compartment. Quantification of the percentage of *Kiss1*-positive neurons that show dendritic expression avoids a possible selection bias arising from the highly heterogeneous distribution of the neurons in this region.

Arc kisspeptin neurons (KP^Arc^) mediate negative feedback regulation of GnRH secretion by estrogen. High estrogen levels suppress *Kiss1* expression in these neurons, and downstream stimulation of GnRH neurons, allowing LH pulsatility and amplitude necessary for normal reproductive function ^(64, 69-71)^. In the arcuate hypothalamic group, the KP neurons co-express *Slc17a6, Esr1, Ar, Pdyn, Tac2*, and have been extensively characterized as “KNDy” neurons ^(72)^. In this study we report that the percentage of *Kiss1*-positive neurons expressing *Kiss1* mRNA in the dendritic compartment was three times higher in males (20.4%) than in proestrous females (6.4%). It has been classically considered that the plasma concentration of estradiol is highest during proestrus ^(73)^; however a recent study in mice showed that although in proestrus the estradiol levels do have a peak, it is during diestrus that estradiol concentrations reach their maximum ^(74)^. Thus, in addition to regulation of dendritic *Kiss1* mRNA in males compared to females, there may also be differences during the estrous cycle in females as well. GNX caused a robust increase in dendritic mRNA labeling within *Kiss1-*expressing cells, of 70% in males and of 79% in females. Thus *Kiss1* dendritic expression in both KP^RP3V^ and KP^Arc^ represents a new arena of KP cellular physiology exploration, albeit requiring correlation with sites of expression in, and release from, these neurons.

While the majority of studies using in situ hybridization report a greater number of KP neurons in the RP3V of female rodents, the data on sex differences in mRNA expression within the ARC remain inconclusive. Some studies report no differences in expression ^(24, 67)^, while others indicate a higher number of *Kiss1*-expressing neurons in males ^(20, 23)^ or in females ^(68)^. These discrepancies may stem from factors such as threshold criteria for *Kiss1*-positivity, the dense neuronal packing within the region, and the varied distribution along the rostrocaudal or mediolateral axes. Focusing on the percentage of *Kiss1* neurons that exhibit dendritic mRNA expression may provide a more consistent and sensitive indicator of expression changes, capturing dynamic cellular responses with greater precision, in neurons with abundant expression of *Kiss1* mRNA.

A third group of KP neurons in the dorsomedial hypothalamus (DMH) was identified and characterized at the mRNA level in the present report. Previous immunohistochemical studies ^(22, 58, 75)^ and Kiss1-CRE mice ^(76, 77)^ showed the prescence of neurons in DMH. However, other studies mapping *Kiss1* expression by in situ hybridization failed to show *Kiss1* mRNA-positive cells in this region ^(6, 20)^. It has been suggested that this inconsistency between immunohistochemical and in situ hybridization histochemical findings might be explained by the detection of Arg-Phe-related peptide-1 (RFRP-1), rather than kisspeptin, using antibodies cross-reacting with both peptides ^(69)^. However, we confirm here the existence of these dispersed KPergic neurons by RNAscope *in situ* hybridization. Since the mRNA probe targets a length of 480 base pairs, almost the entire length of the *Kiss1* mRNA, and is intolerant of partial sequence homology for effective hybridization, this technique confirms the Kiss1+ chemotype of these neurons. Here we characterized these KP-expressing neurons as capable of co-expressing *Slc17a6* (Vglut2 mRNA), *Slc32a1* (VGAT mRNA) and *Ar* (androgen receptor mRNA). No significant changes were detected after gonadectomy in *Kiss1* mRNA expression in this population.

In the ventral premammillary nucleus (PMv) *Kiss1* mRNA puncta were distributed homogeneously among the densely packed cells, seen clearly in parasagittal sections. The existence of this group of KP neurons was suggested earlier by immunohistochemistry in the horse ^(78)^ and indirectly by GFP expression in *Kiss1*-GFP transgenic mice ^(79)^. The authors of the study in the horse interpreted this population as an extension from the Arc neurons, while the results obtained in the transgenic mice study reported expression in several other regions not previously associated with *Kiss1* expression, suggesting ectopic GPF expression. Here, we confirm by *in situ* hybridization the existence of this population. Furthermore, these neurons do not share the molecular signature of the KP^Arc^ population. Apart from expressing *Kiss1* and *Slc17a6*, the KP^PMv^ neurons do not express *Tac2* nor *Pdyn*, and express *Esr1* only weakly. KP^PMv^ neurons also co-express *Adcyap1* and *Nts*, the mRNAs encoding the pituitary adenylate cyclase-activating polypeptide (PACAP) and neurotensin. Interestingly, the Allen Mouse Brain Connectivity Atlas contains documentation, via AAV-aided tracing of genetically targeted PMv Adcyap1-Cre neurons, that these project to the hippocampal CA2 region, a brain region relevant for social behavior ^(80)^.

The extrahypothalamic populations of KP neurons may play distinct physiological and behavioral roles beyond those associated with hypothalamic ones. The KP neurons in the MePD have been studied most extensively ^(17, 24, 55, 57, 75, 81)^. This region is involved in processing olfactory stimuli and influencing matng behavior and reproductive functions ^(82)^. KP^MePD^ neurons co-express vesicular GABA transporter (VGAT) and, to a lesser extent, vesicular glutamate transporter 2 (VGLUT2), indicating a predominantly GABAergic but partly glutamatergic phenotype. This neurotransmitter profile, combined with the expression of androgen receptors (*Ar*) and estrogen receptor alpha (*Esr1*), suggests that MePD KP neurons could influence sociosexual behavior via sex-hormone signaling pathways. The chemoanatomical markers identified in these MePD cells are congruent with a recent study in female mice in which RiboTag/RNA-seq was used to demonstrate Pdyn, Ar, and Esr1 expression within medial amygdala tissue from Kiss1-Cre mice ^(81)^. The male-specific reduction in KP mRNA expression in KP^MePD^ neurons following gonadectomy, reported in this study, supports the putative role of these neurons in male sexual motivation and reproductive behavior. Noteworthy are recent studies using functional magnetic resonance imaging (fMRI), showing that administering KP to healthy men produces enhanced activity in the amygdala and interconected regions such as the hippocampus and cingulate cortex which are crucial to emotional regulation, sexual response and bonding ^(83)^.

The sixth KP population was widely distributed in the BST complex, centered in the septo-hypothalamic nucleus, and with a spatial continuum spanning septal, BST and hypothalamic regions, and is referred to here as KP^SHy^. This KP population shares a molecular signature with KP^MePD^ neurons, expressing both VGAT and sex steroid receptors (Ar and Esr1), suggesting a related functional role in modulating reproductive behaviors. Considering the role of the septum and BST in emotional processing, stress response, motivation and reward-seeking behavior, as well as their connections with the hypothalamus and other limbic structures ^(84-87)^, it is plausible that KP neurons in these regions contribute to modulating these functions. For example, they might influence anxiety levels, motivation related to social interaction, or the integration of emotional states with reproductive behaviors. The similarity in molecular markers and gonadal hormone sensitivity between the MePD and SHy KP neurons highlights a potential role for these neurons in coordinating behaviors that are influenced by both hormonal state and social context, especially in males. We found that in KP^MePD^ and KP^SHy^, gonadectomy induced a decrease in the average number of *Kiss1* mRNA puncta per cell in a sexally dimorphic manner, i.e. only in male mice.

A seventh population of extrahypothalamic KP neurons was found in the brain’s major sensory hub in the dorsal medulla, the solitary tract nucleus (KP^NTS^). KP^NTS^ neurons are glutamatergic and co-express *Tac2, AR*, and *Esr1*, yet do not exhibit changes in *Kiss1* mRNA expression following gonadectomy in either males or females. Our results confirm by ISH the existence of this group of neurons as reported by immunohistochemistry against metastin ^(58, 59)^. As the NTS is a primary sensory and autonomic hub involved in processing visceral and sensory information, KP neurons here may play a role in integrating autonomic signals with overall response to stress or energy demands, rather than reproductive behavior *per se*.

Together, these findings suggest that extrahypothalamic KP neurons, while sharing some common molecular markers, may have region-specific roles that reflect the functional demands of their respective brain areas. KP neurons in the MePD and SHy appear to be closely linked with the modulation of sociosexual and reproductive behaviors, while KP neurons in the NTS might be involved in integrating sensory and autonomic inputs. Further research into the connectivity, physiological responses, and behavioral impact of these KP neurons could provide new insights into the pleiotropic roles of KP signaling in brain function beyond hypothalamic reproductive control.

### 4.2 Distinct responsivities to gonadectomy of the seven KPergic populations

Systematic examination of steroid hormone receptors in both the well-studied hypothalamic and amygdala KP cell groups, and the less-studied hypothalamic and extrahypothalamic groups provide a molecular explanation for the distinction among KP neuronal populations with respect to changes in KP expression upon gonadectomy. Four patterns of alteration to gonadectomy were observed in this study. Classical down-regulation of *Kiss1* expression by GNX in both sexes in RP3V is consistent with high levels of *Esr1* and *Ar* in RP3V, and positive regulation by sex steroids, in both sexes ^(72)^. Confirmation of up-regulation of KP expression by GNX in arcuate nucleus in both sexes (see above) suggests, given high levels of *Esr1* but only low levels of *Ar* in these neurons, that negative modulation of KP expression may occur via estrogen, implying that aromatase expression in nearby neurons may occur in mammals as it does more prominently in seasonally-breeding animals ^(88)^. The third pattern of alteration by gonadectomy was down-regulation in males, but no significant effect in females, in MEA and SHy, again consistent with the relatively high expression of *Ar* but not *Esr1* in these KP cell clusters. The DMH, PMv and NTS showed no significant changes in *Kiss1* expression regulation by GNX in either sex. This finding is consistent with the very low or undetectable expression of either *Esr1* or *Ar* in DMH and PMv, and the lack of *Esr1* in NTS where *Kiss1*-positive cell numbers are low and the expression pattern is sparse.

It is intriguing that of the two major KP cell groups in the hypothalamus, those in the RP3V and in the Arc are oppositely regulated by gonadectomy, and are of opposite chemotype (GABAergic and glutamatergic, respectively). This is consistent with the opposing effects of these two KP populations on reproductive function mediated via GnRH neurons. It has been reported that kisspeptin fibers from these two major KP groups innervate different subpopulations of GnRH neurons, and that there are complex interconnections between these two populations, suggesting a sophisticated feedback loop regulating their activity and ultimatedly impacting GnRH function ^(89)^. A recent study using electrophysiological recordings in combination with optogenetics, showed excitatory and inhibitory projections from the KP^Arc^ and the KP^RP3V^ respectively, converging onto preautonomic paraventricular (PVN) and dorsomedial (DMH) neurons to control their excitability ^(90)^. How these post-synaptic effects are modulated by co-release of KP, glutamate and GABA from KP^RP3V^ and KP^Arc^ neurons is a question awaiting more detailed investigation.

Significant advance in our understanding of the key role of kisspeptin in human sexually-related behavior has been provided by clinical studies of kisspeptin administration in men and women ^(83, 91)^. However, except for a few reports (e.g. Pineda et al. 2017 ^(17)^), there has been little exploration of kisspeptin projections to areas other than GnRH neurons of the hypothalamus. As pointed out by Liu and Herbison ^(41)^, the post-synaptic effects of kisspeptin leading to behavioral effects such as increased sexual responsiveness may be mediated via neuromodulatory rather than direct neurotransmitter mechanisms, and at sites of kisspeptinergic innervation of brain areas involved in both sensorial processing and behavioral state control. Thus, a better understanding of the projection of KP neurons outside of the hypothalamus, arising from both hypothalamic and extrahypothalamic KP cell bodies, is needed. The chemotyping of KP neurons provided here for the mouse provides a context for consideration of how different sets of KP neurons in the mammalian brain work together to orchestrate male and female reproduction, not only via regulation of individual endocrine responses, but by regulation of individual and social behaviors.

## Acknowledgements

The research leading to these results received funding from the National Autonomous University of Mexico, under Grant Agreement UNAM-PAPIIT-IG200121 (LZ), IA202724 (ORH-P), Mexican Nacional Council for Humanity, Science and Technology (CONAHCYT), under grant agreement CF-2023-G-243 (LZ), and NIMH-IRP, NIH, under Grant Agreement MH002386 (LEE). The investigators of this study have been supported by the following fellowships: sabbatical year fellowship from the PASPA program of the Dirección General de Personal Académico (DGAPA) of the Universidad Nacional Autónoma de México (UNAM) (LZ, VSH); Mexican CONAHCYT sabbatical fellowship (LZ); Fulbright-García Robles Fellowship (VSH); DGAPA-UNAM POSDOC program (MAZ) and PREI program for a sabbatical research stay (RPM) in LZ’s lab in UNAM, Mexico. RPM thanks the University of Pretoria for sabbatical leave and the National Research Foundation and Medical Research Council of South Africal for research grant support.

